# SSTR2-targeted theranostics in hepatocellular carcinoma

**DOI:** 10.1101/2024.10.16.618740

**Authors:** Majid Momeny, Solmaz AghaAmiri, Servando Hernandez Vargas, Belkacem Acidi, Sukhen C. Ghosh, Tyler Bateman, Jack Adams, Vahid Khalaj, Ahmed O. Kaseb, Hop S. Tran Cao, Ali Azhdarinia

## Abstract

**Background:** While the clinical use of radiolabeled somatostatin analogs is established in neuroendocrine tumors, there is significant interest in expanding their use for other somatostatin receptor 2 (SSTR2)-expressing cancers. This study investigates the utility of SSTR2-targeted theranostics in hepatocellular carcinoma (HCC);

**Methods:** We measured SSTR2 expression in HCC cell lines and clinical samples using qRT-PCR, Western blot, and a public dataset. We evaluated [^67^Gallium]Ga-DOTATATE uptake, tested [^177^Lutetium]Lu-DOTATATE cytotoxicity, and assessed [^68^Gallium]Ga-DOTATATE tumor targeting in HCC animal models and a patient via PET/CT;

**Results:** SSTR2 expression was confirmed in HCC cell lines and clinical samples. Radioligand uptake studies validated SSTR2-mediated [^67^Gallium]Ga-DOTATATE uptake, and [^177^Lutetium]Lu-DOTATATE treatment reduced cell proliferation. [^68^Gallium]Ga-DOTATATE PET/CT scans detected tumors in animal models and spinal metastases in a patient with HCC;

**Conclusion:** These findings suggest for the first time that SSTR2-based theranostics could have strong implications for detection and treatment of HCC.

**Simple Summary:** This study investigates the use of SSTR2-targeted theranostics, combining diagnostic and therapeutic approaches, in hepatocellular carcinoma (HCC). We confirmed significant SSTR2 expression in HCC cells and patient samples, showing that radiolabeled compounds such as [^67^Ga]Ga-DOTATATE and [^177^Lu]Lu-DOTATATE, commonly used in neuroendocrine tumors, could also target HCC. In preclinical models and a patient case, PET/CT imaging and treatments demonstrated effective tumor detection and shrinkage. These findings suggest that SSTR2-targeted theranostics could offer a novel, targeted method for diagnosing and treating HCC, potentially improving outcomes for patients with this challenging cancer.

## 1. Introduction

Somatostatin receptor 2 (SSTR2) is a G-protein-coupled receptor that regulates key endocrine and nervous system functions by mediating the inhibitory effects of somatostatin on hormone secretion, neurotransmitter release, and cell proliferation. Under normal physiological conditions, SSTR2 is expressed in tissues such as the brain, gastrointestinal tract, and pancreas. However, in neuroendocrine tumors (NETs), SSTR2 is often significantly overexpressed, making it a highly attractive target for both diagnostic imaging and peptide receptor radionuclide therapy (PRRT) through theranostic approaches. Theranostics, which combines diagnostic precision with therapeutic efficacy, facilitates early disease detection, targeted drug delivery, and minimizes damage to healthy tissues [1].

The overexpression of SSTR2 in NETs has enabled the development and FDA approval of radiolabeled somatostatin analogs, such as [^68^Gallium]Ga-DOTATATE for PET imaging and [^177^Lutetium]Lu-DOTATATE for PRRT, offering precise tumor visualization and effective targeted therapy [2,3]. Clinical studies have shown that SSTR2-targeted theranostics significantly improve clinical outcomes, with radiolabeled DOTATATE analogs reducing disease progression and mortality in NET patients by approximately 80% [4]. These promising results underscore the potential of SSTR2-based theranostics and support the exploration of similar approaches in other SSTR2-overexpressing malignancies.

Liver cancer is the fourth leading cause of cancer death and the sixth most diagnosed cancer globally, with over one million cases projected by 2025 [5]. In the U.S., liver cancer death rates rose by 43% from 2000 to 2016 [6]. Hepatocellular carcinoma (HCC), the most common liver cancer, has a 5-year survival rate of 18%, making it the second deadliest cancer after pancreatic cancer [6,7]. Treatment outcomes for advanced HCC remain poor, with limited benefits from conventional therapies and systemic agents like sorafenib [8-10]. Immunotherapies like atezolizumab are now frontline treatments [11], but drug resistance, high recurrence rates, and lack of early detection biomarkers highlight the need for better strategies [12].

Studies show that about 40% of HCC samples have positive SSTR2 membrane staining: 9.6% strong, 21.2% moderate, and 7.7% weak [13]. A clinical trial found that HCC patients with positive [^68^Galluim]Ga-DOTATATE PET/CT had better survival outcomes when treated with a somatostatin analog, indicating a subset of HCC patients could benefit from SSTR2-targeted theranostics [14]. These findings provide a strong rationale to investigate SSTR2-targeted theranostics in HCC.

## 2. Materials and Methods

### Antibodies and chemicals

SSTR2 (clone A-8) and β-actin (clone C4) were purchased from Santa Cruz Biotechnology. [^177^Lutetium]LuCl3 from the National Isotope Development Center was used to produce [^177^Lutetium]Lu-DOTATATE via standard procedures [15]. DOTATATE was radiolabeled with [^68^Gallium]GaCl3 (RLS Radiopharmacies) or [^67^Gallium]-GaCl3 (Cardinal Health) following established methods [16]. Radiochemical yields and purity were assessed by instant thin-layer chromatography and radio-HPLC, and the final products were diluted with PBS as needed.

### Cell culture

The SNU449, Hep3B2, HepG2, and H69 cell lines were obtained from the American Type Culture Collection, while Huh7 cells were sourced from the Japanese Collection of Research Bioresources Cell Bank. BON-1 and QGP-1 cells were provided by Dr. Jeffry A. Frost, The University of Texas Health Science Center Houston. The cells were cultured according to the recommendations provided by their respective sources.

### qRT-PCR, Western blotting, and WST-1 cell proliferation assay

These procedures followed previous descriptions [17]. For qRT-PCR, the primers used were: SSTR2 (F: GAGGAGCCAGGAACCCCAAA, R: GGATCCAGTGTGACATCTTTGCT) and GAPDH (F: TCGGAGTCAACGGATTTGGTC, R: TGAAGGGGTCATTGATGGCA).

### cBioPortal and the Human Protein Atlas Database

Genetic mutations and mRNA expression levels of SSTR2 in HCC patients were analyzed using the TCGA HCC dataset [18]. The Human Protein Atlas (https://www.proteinatlas.org/) was used to investigate SSTR2 protein expression across various cancers and to examine immunohistochemistry (IHC) slides for SSTR2 expression in HCC patients.

### Uptake study

For radioactive uptake studies, 2 × 10^5^ cells in 96-well plates were incubated with 2 nM [^67^Gallium]Ga-DOTATATE, with or without 100X octreotide block, for 1 h at 37 °C. After removing unbound radioligand, cell-associated radioactivity was quantified with a gamma counter, and the percentage of total radioactivity was calculated from a known aliquot.

### Animal studies

All procedures followed ethical guidelines approved by the IACUC at The University of Texas Health Science Center at Houston. Subcutaneous tumor models were created by injecting 3 × 10^6^ Huh7 or SNU449 cells into the left shoulder of athymic nude mice (n=5/group). HCT116 colon cancer cells, which do not express SSTR2 [19], served as negative controls. Mice received an intravenous injection of 7.4 MBq (200 μCi, 2 nmol) of [^68^Gallium]Ga-DOTATATE and underwent PET/CT imaging (Bruker Albira) 1 h later. Tissues were then harvested for ex vivo biodistribution and radioactivity measurement.

### Immunohistochemistry (IHC)

Tumor tissues were fixed, paraffin-embedded, sectioned, and stained with H&E. For IHC, sections underwent antigen retrieval, then were incubated overnight with anti-SSTR2 antibody (ab134152, Abcam) and followed by a biotinylated goat anti-rabbit IgG secondary antibody. Detection was carried out using a DAB kit (ab64261, Abcam).

### Statistical analysis

Data were graphed and analyzed using GraphPad Prism Software 10.2.3 using two-way ANOVA followed by Šídák’s multiple comparisons test. The data are presented as mean ± standard deviation (SD).

## 3. Results

Analysis of the TCGA HCC dataset [18] revealed SSTR2 amplification or overexpression in a significant proportion of HCC patients (Fig. 1A). The Human Protein Atlas dataset showed SSTR2 protein expression in most HCC patients (Fig. 1B, C). We measured SSTR2 expression in HCC cell lines SNU449, Hep3B2, Huh7, and HepG2 using qRT-PCR and Western blotting, finding high levels of SSTR2 mRNA and protein (Fig. 1D, E). Compared to NET cell lines H69 and QGP1, HCC cell lines had significant *SSTR2* mRNA levels (Fig. 1D). Radioligand uptake studies confirmed substantial SSTR2-mediated [^67^Gallium]Ga-DOTATATE uptake in SNU449, Huh7, and HepG2 cells, comparable to NET cell lines H69 and BON-1 (Fig. 1F).

**Figure 1.**
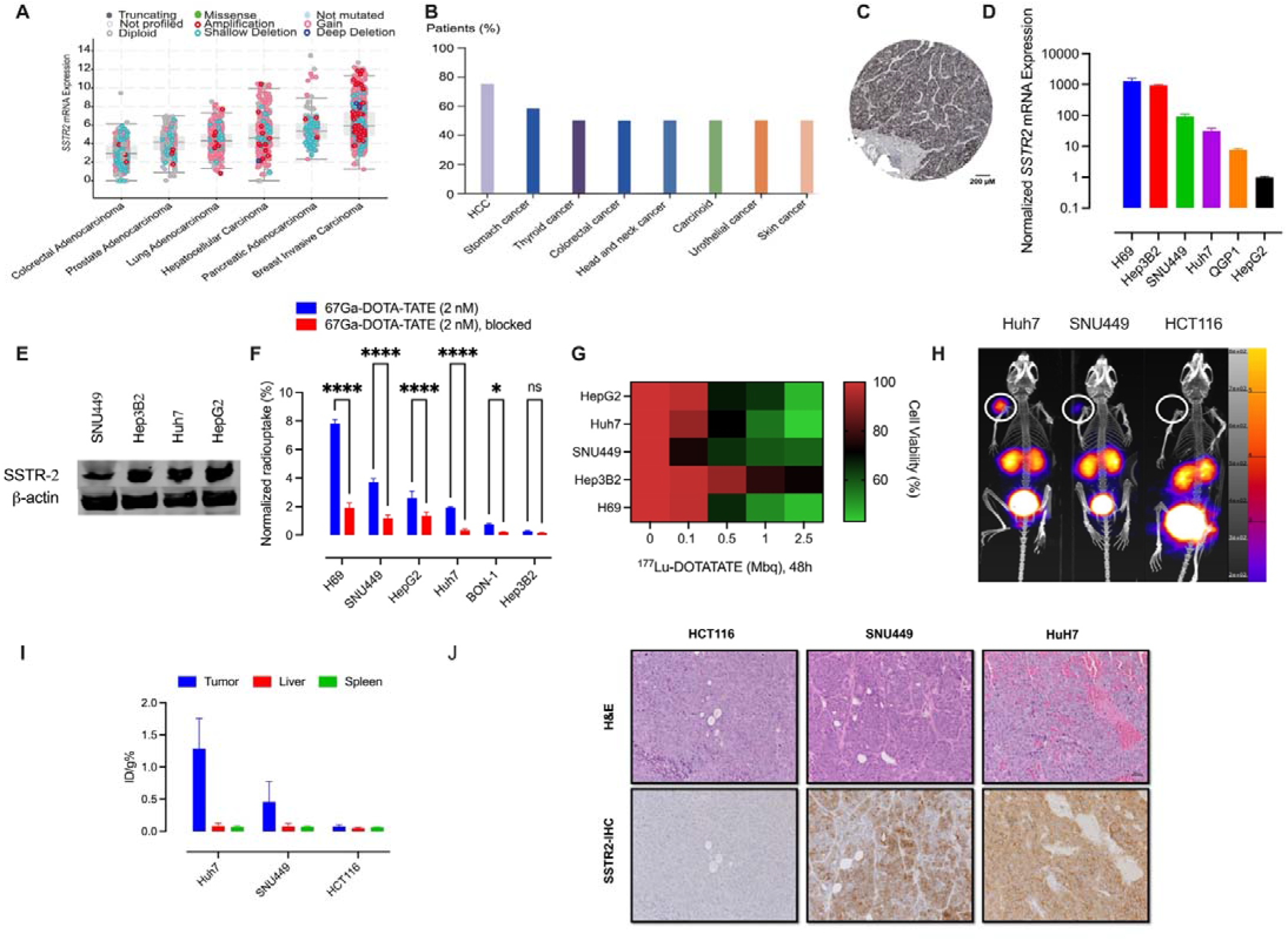
(A) SSTR2 is amplified and/or overexpressed in a considerable percentage of HCC patients in the TCGA hepatocellular carcinoma dataset [18]. (B) SSTR2 protein expression in HCC patients, as compared with other solid tumors. The image is extracted from the Human Protein Atlas dataset (https://www.proteinatlas.org). (C) IHC staining for SSTR2 in an HCC patient (patient id #2766) was extracted from the Human Protein Atlas dataset. The anti-SSTR2 antibody #HPA007264 from Sigma was used for staining. (D) The mRNA levels of SSTR2 were determined by qRT-PCR. The data were evaluated in triplicate and collected from three independent experiments. Gene expression levels were normalized to *GAPDH* in each cell line and then normalized to HepG2 cells. (E) SSTR2 protein expression in the HCC cell lines was examined by Western blot analysis and compared with NCI-H69 small cell lung cancer cells. β-actin was used as the loading control. (F) Uptake of [^67^Gallium]Ga-DOTATATE in HCC cells as compared with NCI-H69 and BON-1 cells. Octreotide (100X) was used to block the receptor. Data were analyzed by two-way ANOVA followed by Šídák’s multiple comparisons test. Statistically significant values of *p < 0.05 and ****p < 0.0001 were determined. (G) The anti-proliferative activity of [^177^Lutetium]Lu-DOTATATE in the HCC cell lines as measured by WST-1 cell proliferation assay (Sigma). NCI-H69 cells were used as a benchmark. The data were evaluated in triplicate and collected from three independent experiments. (H) Maximum-intensity projection showing tumor-specific [^68^Gallium]Ga-DOTATATE uptake in subcutaneous models of Huh7 and SNU449. Mice were i.v. injected with 7.4 MBq of [^68^Gallium]Ga-DOTATATE into the tail vein and PET/CT imaging was performed at 1 h p.i. (I) Quantification of radioactive biodistribution in subcutaneous models of Huh7, SNU449, and HCT116. (J) Corresponding IHC analysis showing SSTR2 expression in FFPE tumor sections from the mice with the HCC tumors. The tumor from the HCT116-bearing mice was used as the negative control.

To assess the anti-proliferative effects of [^177^Lutetium]Lu-DOTATATE, HCC cell lines were treated with increasing radioactivity for 48 h. SNU449, Huh7, and HepG2 cells showed a significant decrease in viability, while Hep3B2 cells showed minimal effects (Fig. 1G). This result, consistent with radioligand uptake profiles, suggests that NET-based theranostic approaches (e.g., H69) may also be effective in HCC cells.

To evaluate SSTR2-targeted imaging in HCC models *in vivo*, we used subcutaneous mouse models with Huh7 and SNU449 cells, and HCT116 cells as a negative control. After IV injection of [^68^Gallium]Ga-DOTATATE and PET/CT imaging, both HCC tumors were clearly visible, with radiotracer accumulation correlating with SSTR2 expression levels (Fig. 1H). *Ex vivo* biodistribution analysis confirmed PET results, showing a 16.7-fold higher [^68^Gallium]Ga-DOTATATE uptake in Huh7 cells compared to controls (Fig. 1I).

IHC analysis confirmed SSTR2 expression in tumor sections from HCC xenografts along with the absence of the target in HCT116 tumors (Fig. 1J). Notably, a [^68^Gallium]Ga-DOTATATE PET/CT scan conducted on a patient with a history of HCC demonstrated SSTR2 positivity in spinal metastases (Fig. 2). The patient had previously undergone definitive radiation therapy for HCC and later required a small bowel resection. During routine surveillance for NET, a [^68^Gallium]Ga-DOTATATE PET/CT scan detected a tumor in the spine. Initially suspected to be a metastasis from the NET, biopsy of the lesion confirmed it was HCC. Collectively, these novel findings illustrate the roles of SSTR2 as a functional biomarker for theranostic-based detection and treatment in HCC and emphasize the need for further clinical investigations.

**Figure 2.**
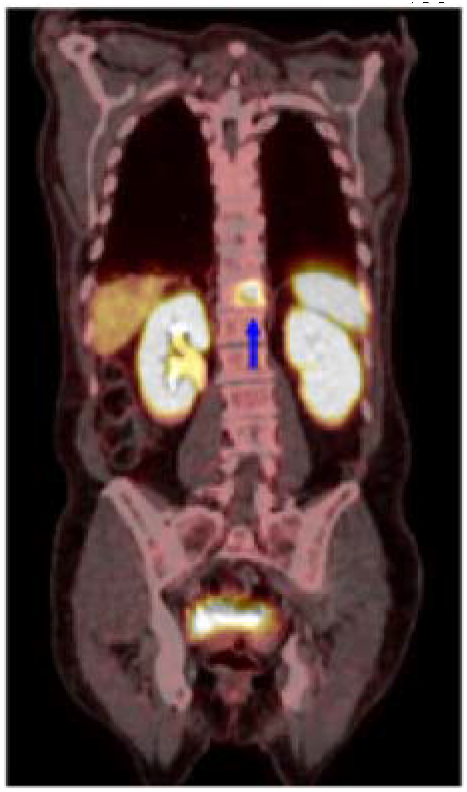
A [^68^Gallium]Ga-DOTATATE PET/CT scan showing SSTR2 positivity in spinal metastases from a patient with HCC.

## 4. Discussion

In this study, we investigated the utility of SSTR2-targeted theranostics in the context of HCC. Examination of SSTR2 expression in HCC cell lines and clinical specimens revealed notable uptake of [^67^Gallium]Ga-DOTATATE in 75% of examined HCC cell lines. Subsequent treatment with [^177^Lutetium]Lu-DOTATATE resulted in a significant reduction in cell viability, suggesting potential utility of this approach for treating HCC. We also showed that [^68^Gallium]Ga-DOTATATE PET/CT imaging can effectively localize HCC in subcutaneous tumor models with varying SSTR2 expression levels and an HCC patient with spinal metastases. This observation, in combination with the low background signal from PET and *ex vivo* biodistribution analysis, indicates the feasibility of tumor delineation in patients with HCC. Our results present compelling evidence for the feasibility and the potential diagnostic visualization and therapeutic impact of SSTR2-targeted theranostics in the management of HCC, warranting further clinical exploration.

## 5. Conclusions

In conclusion, our study demonstrates the potential of SSTR2-targeted theranostics for HCC. We observed significant [^67^Gallium]Ga-DOTATATE uptake in 75% of HCC cell lines and reduced cell viability with [^177^Lutetium]Lu-DOTATATE treatment. Additionally, [^68^Gallium]Ga-DOTATATE PET/CT successfully localized tumors in models with different SSTR2 expression and an HCC patient. These findings support further clinical exploration of SSTR2 theranostics in HCC management.

## Author Contributions

All authors contributed to the study conception and design. Material preparation, data collection and analysis were performed by Majid Momeny, Solmaz AghaAmiri, Servando Hernandez Vargas, Belkacem Acidi, Sukhen C. Ghosh, Tyler Bateman, Jack Adams, Vahid Khalaj, Ahmed O. Kaseb and Hop S. Tran Cao. The first draft of the manuscript was written by Majid Momeny and all authors commented on previous versions of the manuscript. The study was supervised by Majid Momeny and Ali Azhdarinia. All authors read and approved the final manuscript.

## Funding

This work was supported by the John S. Dunn Research Scholar Fund.

## Institutional Review Board Statement

All animal studies were performed in accordance with ethical protocols approved by the Institutional Animal Care and Use Committee (IACUC) of The University of Texas Health Science Center at Houston.

## Informed Consent Statement

“Informed consent was obtained from all subjects involved in the study.”

## Data Availability Statement

Data from this study are available upon reasonable request from the corresponding authors.

## Acknowledgments

We would like to thank Dr. Jeffry A. Frost from The University of Texas Health Science Center at Houston for generously providing the BON-1 and QGP-1 cell lines used in this study.

## Conflicts of Interest

“The authors declare no conflict of interest.”

## Notes

### Competing Interest Statement

The authors have declared no competing interest.

